# Aberrant Epithelial Differentiation Contributes to Pathogenesis in a Murine Model of Congenital Tufting Enteropathy

**DOI:** 10.1101/2020.10.12.330522

**Authors:** Barun Das, Kevin Okamoto, John Rabalais, Jocelyn Young, Kim E. Barrett, Mamata Sivagnanam

**Author notes:** Corresponding Author: Mamata Sivagnanam, Division of Gastroenterology, Hepatology and Nutrition, Department of Pediatrics, 9500 Gilman Drive, La Jolla, CA 92093 USA, University of California San Diego, Rady Children’s Hospital San Diego.

## Abstract

**Background & Aims:** Congenital Tufting Enteropathy (CTE) is an intractable diarrheal disease of infancy caused by mutation of Epithelial Cell Adhesion Molecule (EpCAM). The cellular and molecular basis of CTE pathology has been elusive. We hypothesized that the loss of EpCAM in CTE results in altered lineage differentiation and defects in absorptive enterocytes thereby contributing to CTE pathogenesis.

**Methods:** Intestine from CTE mice was evaluated for specific markers by RT-qPCR, western blotting and immunostaining. Body weight, blood glucose and intestinal enzyme activity were also investigated. A CTE enteroid model was used to assess whether the decreased census of secretory cells could be rescued.

**Results:** CTE mice exhibited alterations in brush-border function, disaccharidase activity and glucose absorption, potentially contributing to nutrient malabsorption and impaired weight gain. Altered cell differentiation in CTE mice led to decreased secretory cells and increased numbers of absorptive cells, though the absorptive enterocytes lacked key features, causing brush border malfunction. Further, treatment with Notch signaling inhibitor, DAPT, increased the numbers of major secretory cell types in CTE enteroids (Graphical abstract 1).

**Conclusions:** Alterations in intestinal epithelial cell differentiation in CTE mice favor an increase in absorptive cells at the expense of secretory cells. Although the proportion of absorptive enterocytes is increased, they lack key functional properties. We conclude that these effects underlie pathogenic features of CTE such as malabsorption and diarrhea, and ultimately the failure to thrive seen in patients. The ability of DAPT to reverse aberrant differentiation suggests a possible therapeutic strategy.

**Synopsis:** A murine model of Congenital Tufting Enteropathy exhibits altered intestinal cell differentiation, leading to increased absorptive and decreased secretory cells, which can be reversed with DAPT. Absorptive enterocytes in these mice are also dysfunctional, contributing to disease pathogenesis.

**Graphical Abstract:** 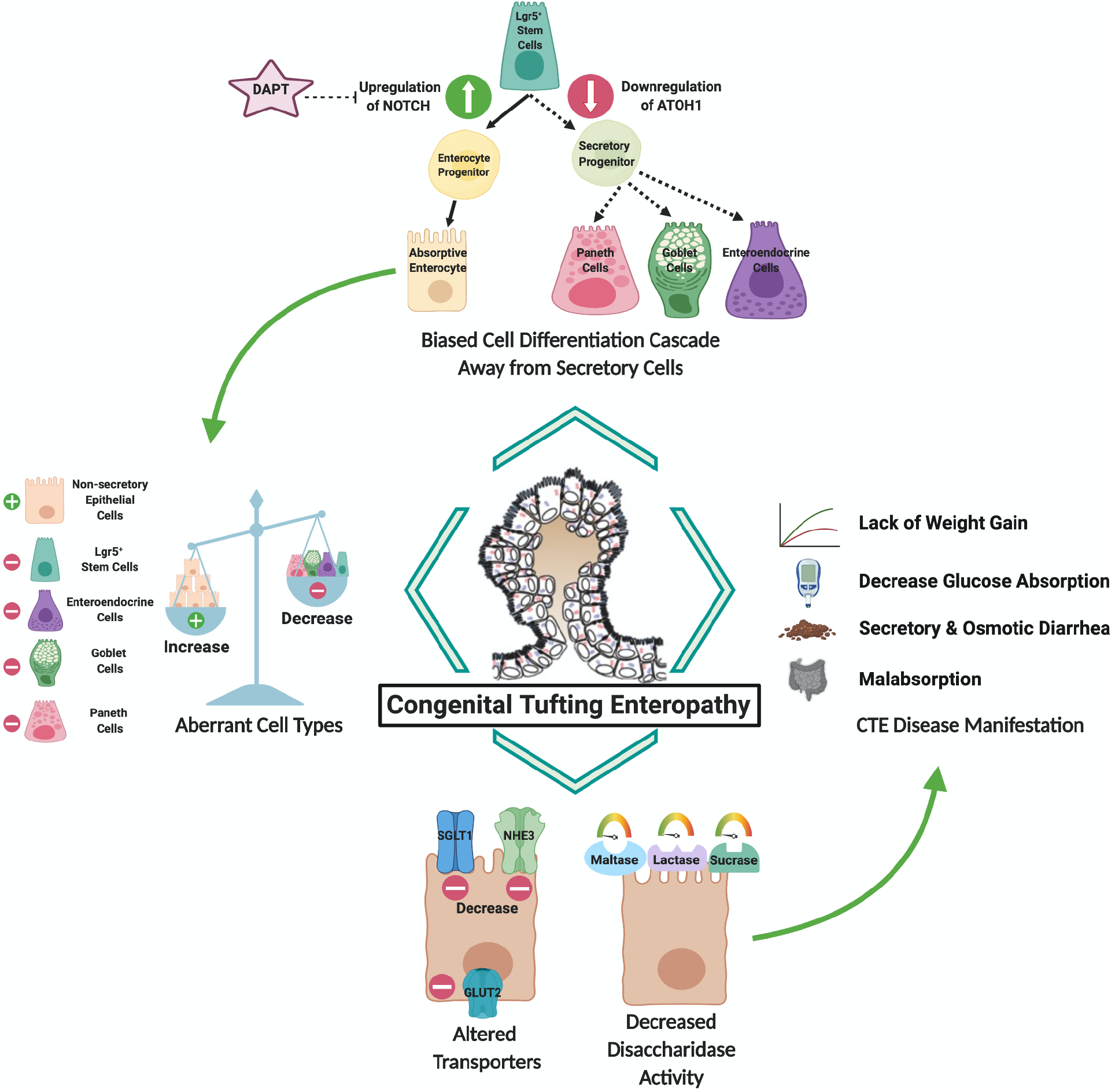

## Introduction

Congenital Tufting Enteropathy (CTE) [Online Mendelian Inheritance in Man (OMIM): 613217], is a rare intractable diarrheal disease of infancy characterized by profuse watery diarrhea, electrolyte imbalances, and impaired growth [1, 2]. Its intestinal pathology includes villous atrophy, crypt hyperplasia, and epithelial tufts leading to intestinal failure. CTE belongs to a family of congenital diarrhea and enteropathies that are both secretory and osmotic in nature [1]. Mutations in Epithelial Cell Adhesion Molecule (EpCAM) were identified as the primary cause for CTE [3]. Although Serine Peptidase Inhibitor Kunitz Type 2 (SPINT2) mutations have also been reported in a syndromic form of the disease, most CTE patients possess mutations in EpCAM [4]. Patients with CTE typically depend on parenteral nutrition and rarely achieve enteral autonomy without intestinal transplant. The significant morbidity and mortality associated with the disease suggest an urgent need for an improved understanding of disease pathophysiology, with the hope of contributing to the development of therapeutics.

Studies utilizing various *in vivo*, *ex vivo* and *in vitro* generated models of CTE with EpCAM mutations have revealed some insights regarding the onset of the disease phenotype. EpCAM knockout mice have dysregulated E-cadherin and β-catenin in the enteric mucosa, leading to crypt-villus disorganization [5]. A study using a neonatal murine model bearing an EpCAM mutation identified in a CTE patient showed neonatal lethality, growth retardation with epithelial tufts, enterocyte crowding, altered desmosomes and intercellular gaps similar to human CTE patients [6]. Another study of adult mice bearing the same mutation revealed growth retardation, CTE-like histopathology, impaired intestinal barrier function and decreases in the tight junction proteins zonula occludens-1 (ZO-1) and occludin [7]. CTE intestinal organoids from mice reveal alterations in differentiation in addition to barrier dysfunction [8]. Moreover, data suggest that deficiency of EpCAM in a colonic cell line is accompanied by ion transport and barrier defects [7]. These barrier defects may contribute to the secretory portion of CTE-associated diarrhea [9–11], but do not clearly explain the osmotic aspects. Further, while CTE patients have features of nutrient malabsorption, the role epithelial dysfunction plays in this aspect of the disease has yet to be understood.

The fact that most mice with engineered mutations in EpCAM exhibited intestinal failure suggests an important role for EpCAM in maintaining intestinal epithelial organization [5, 6, 11, 12]. However, there is still much to be elucidated about the mechanistic role of mutant EpCAM in intestinal cellular organization. Via a tightly regulated program of differentiation, stem cells give rise to intestinal epithelial cells (IEC) that can be categorized as either absorptive cells or as four different secretory cell lineages: goblet, Paneth, enteroendocrine and tuft cells. Intestinal stem cells continuously self-renew to generate an intermediate cell type, known as transit amplifying cells, which then undergo terminal differentiation and maturation to absorptive and secretory IEC to maintain intestinal cellular homeostasis [13]. The NOTCH and ATOH1 (MATH1) pathways play a crucial role in intestinal function by regulating the choice between absorptive and secretory lineages [14–17]. Previous studies in CTE patients, CTE mice and CTE enteroids revealed decreases in Paneth and goblet cell numbers [8], which may indicate an alteration in intestinal epithelial cell differentiation. Thus, detailed investigation of the intestinal epithelial cell differentiation pathway and its contributions to CTE pathogenesis was of interest.

Recent studies have also revealed a role for defective absorptive enterocytes in the pathogenesis of various intestinal conditions including congenital diarrheal disorders [18] such as congenital lactase deficiency [19], glucose-galactose malabsorption [20], sucrase-isomaltase deficiency [21], congenital chloride diarrhea [22], and microvillous inclusion disease [23]. Broadly, these diseases are accompanied by deficits in brush border-associated enzymes and transporters, as well as defects in intracellular protein transport, intracellular lipid transport and intestinal barrier function [24]. Past studies utilizing different CTE models primarily focused on intestinal barrier function disruption and tight junction disorganization. However, the impact of loss of EpCAM on absorptive enterocyte maturation or function has not been studied. We reasoned that a deeper knowledge of the differentiation of absorptive enterocytes would broaden our understanding of CTE pathobiology.

In the present study, we hypothesized that a loss of function mutation of EpCAM results in an altered lineage differentiation cascade and defective absorptive enterocytes that contribute to CTE pathogenesis. Hence, the IEC differentiation cascade and functional markers of mature absorptive enterocytes are systematically evaluated.

## Results

### CTE mice exhibit impaired weight gain upon disease onset

To expand our knowledge of primary and ancillary pathological features due to mutation of EpCAM, our previously described adult CTE mice with inducible expression of mutant EpCAM were utilized [7]. While characteristic features of small intestinal histopathology and epithelial barrier properties of CTE had already been described in these mice, failure to thrive or poor weight gain had not been evaluated, despite it being a predominant symptom of CTE [7]. We therefore monitored the growth of both control and mutant mice over the course of 5 days. While control mice exhibited steady weight gain, mutant mice were noted to have impaired growth beginning on day 4 of the observation period (Fig 1A), similar to CTE patients and our previous study in neonatal mice [6].

**Figure 1.**
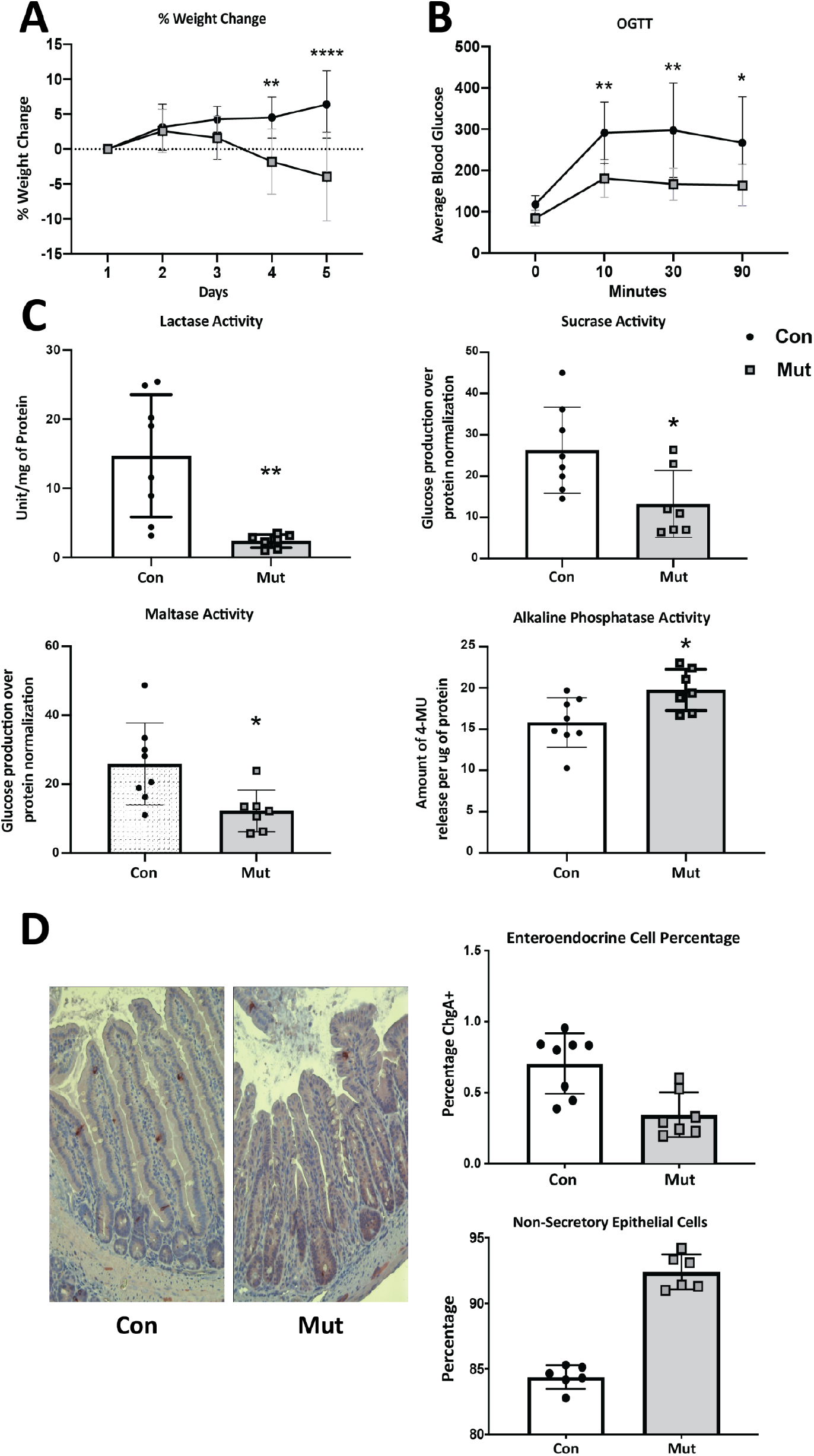
Pathologic features of CTE mice. **(A)** Percentage (%) weight change from day 1 in CTE mice (Mut) and control mice (Con) over 5 days with weight loss in Mut compared to Con mice (n=8). Tamoxifen was administered on day 1 and 2 to induce expression of mutant EpCAM. **(B)** Oral glucose tolerance test at day 5. Decreased blood glucose levels in Mut compared to Con mice at 10, 30, and 90 minutes (n=5). **(C)** Decrease in lactase, sucrase and maltase activity and increased alkaline phosphatase activity in duodenal lysate of Mut (n=7) compared to Con mice (n=8) at day 5 of treatment **(D)** Representative IHC brightfield (20X) images showing decreased percentage of ChgA^+^ cells (enteroendocrine cells) in Mut (n=7) compared to Con mice (n=8) and increased number of non-secretory epithelial cells relative to total epithelial cells (n=6). Scalebar represents 100 μm. *=p<0.05, **=p<0.01, ***=p<0.001, ****=p<0.0001 by Student’s t test in all results.

### Impaired glucose absorption and lack of digestive enzyme activity in CTE mice

The lack of weight gain in mutant mice led us to evaluate nutrient absorption in these animals. Nutrient malabsorption can be attributed to both defective absorption and lack of digestion. As CTE affects the intestinal epithelium that contributes to both of these physiological processes, we set out to investigate whether malabsorption was a consequence of an inappropriate mechanism of absorption and/or to a lack of intestinal brush border hydrolases. Carbohydrates are a primary source of caloric intake, especially in children, thus carbohydrate absorption was examined in the form of an oral glucose tolerance test (OGTT). Glucose uptake was significantly blunted in mutant mice compared to controls at 10, 30, and 90 minutes after glucose administration (Fig 1B).

To investigate whether brush border hydrolases are impacted by CTE in our mouse model, enzymes exclusively synthesized by enterocytes in the small intestine were the primary focus. We measured mucosal activity of sucrase, maltase and lactase in CTE mice. Activity of all three enzymes was significantly decreased in mutant mice compared to controls (Fig 1C). Among the three, lactase activity was most severely affected (average 8 fold). To study whether the decreased hydrolase activity simply reflects the altered intestinal epithelial structure in the mutant mice, we also measured activity of intestinal alkaline phosphatase. In contrast to the other digestive enzymes studied, intestinal alkaline phosphatase activity was elevated in mutant mice compared to controls (Fig 1C). Overall, our data suggest that a significant and selective decrease in brush border digestive enzyme activity contributes to the malabsorption seen in CTE.

### CTE mice have decreased numbers of enteroendocrine cells and increased numbers of absorptive cells

A significant decrease in Paneth and goblet cell numbers in CTE patients and CTE mice has previously been reported [8]. In order to assess whether other secretory cells types are affected in CTE, we evaluated numbers of enteroendocrine cells in the CTE mice. Staining for the specific enteroendocrine cell marker, chromogranin A (CHGA) revealed a decrease in the percentage of enteroendocrine cells in CTE mice compared to controls (Fig 1D). This result confirms that all secretory epithelial cell types are likely affected in CTE. We also examined absorptive cells in the mutant mice by counting all non-secretory epithelial cells in the intestinal epithelium. Despite evidence for decreased epithelial absorption in the CTE mice, the number of absorptive epithelial cells was significantly increased in CTE mice compared to controls (Fig 1D). This is also consistent with our finding of increased alkaline phosphatase activity.

### Evaluation of gene expression characteristic of intestinal epithelial cell types

The apparent reciprocal alteration in secretory/absorptive intestinal cell types in CTE mice led us to further investigate cell differentiation at the molecular level. Since CTE mice have decreased enteroendocrine cells compared to controls, ChgA was evaluated at the gene level. In line with the immunohistochemistry (IHC) findings (Fig 1D), mRNA expression for ChgA was decreased in mutant mice compared to controls (Fig 2A). In order to further understand the dynamics of intestinal epithelial cell differentiation in CTE mice, markers that define intestinal stem cells (ISCs), progenitor cells, transit amplifying cells, and enterocyte differentiation markers were chosen. The classical long-lived, self-renewing multipotent ISCs were assessed by studying expression of leucine-rich repeat-containing G-protein coupled receptor 5 (*LGR5*) and BMI1 proto-oncogene, polycomb ring finger (*BMI1*) [25]. There was a significant decrease in expression of Lgr5, a marker of mitotically active ISCs, in mutant mice compared to controls (Fig 2A). On the contrary, expression of BMI1, a marker of quiescent ISCs, was not changed between CTE mutants and controls. Similarly, there was no difference in mRNA levels for the ISC regulator Achaete-Scute Family BHLH Transcription Factor 2 (ASCL2) [26], the robust surrogate stem cell marker Olfactomedin-4 (*OLFM4*)[27], or the early lineage/transit amplifying progenitor/stem cell markers Prominin 1 (*PROM1*) [28], Musashi RNA Binding Protein 1 (MSI1) [29], and intestinal epithelial differentiation homeobox transcription factor Caudal Type Homeobox 2 (CDX2) [30] between control and mutant mice (Fig 2A). These results indicate there is decreased mRNA expression for markers of some cell types (enteroendocrine cells and ISC) in CTE mice while markers for other stem or progenitor cells and differentiation markers are unchanged., indicating no direct role of EpCAM in those cell types.

**Figure 2.**
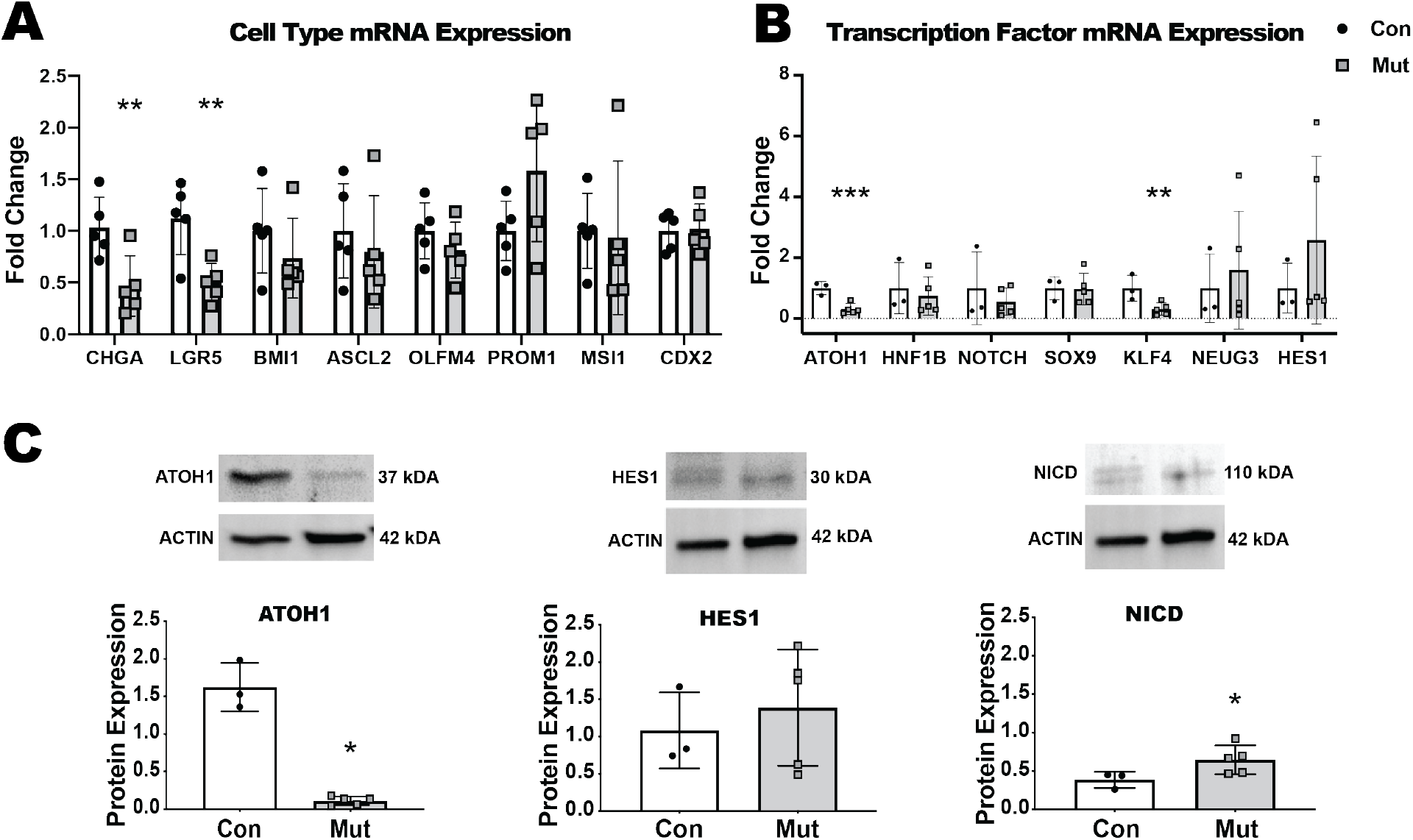
Intestinal epithelial cell differentiation in CTE mice. **(A)** Cell type mRNA expression. Decreased expression of Chromogranin A (CHGA, enteroendocrine cell) and LGR5; With normal levels of progenitor, and transient amplifying cell markers *BMI1, ASCL2, OLFM4, PROM1, MSI1*, and *CDX2* in small intestinal tissue lysates from CTE mice (Mut) compared to control mice (Con) (n=5). **(B)** mRNA expression of transcription factors. Decreased expression of *ATOH1*, and *KLF4* with stable expression of *HNF1b*, *NOTCH*, *SOX9*, *NEUG3*, and *HES1* in small intestinal tissue lysates of Mut compared to Con mice (n=5). **(C)** Representative western blot images for ATOH1, HES1, and NICD relative to actin in small intestinal tissue lysates from Con and Mut mice (n=5). Decreased protein expression of ATOH1 with unchanged expression of HES1 and increased NICD in Mut compared to Con mice. Same blot was used to detect HES1 and NICD protein. *=p<0.05, **=p<0.01, ***=p<0.001, ****=p<0.0001 by Student’s t test for all results.

### CTE mice show alterations in intestinal stem cell differentiation cascades

Our finding of a decreased number of secretory IECs along with decreased gene expression of ChgA and Lgr5 in CTE mice led us to study signaling molecules and transcription factors that regulate the secretory pathway and their role in IEC differentiation. ATOH1 has been shown to be essential for secretory cell differentiation both in the adult intestine and in the embryo [13]. Thus, cells expressing ATOH1 are driven toward a secretory cell lineage (Paneth, enteroendocrine, goblet) compared with those driven by Notch, which instead follow an absorptive path [31, 32]. A significant decrease in mRNA for *ATOH1* in CTE mice corresponded with the observed decrease in secretory IEC lineages (Fig 2B). This was further confirmed by detecting decreased ATOH1 protein in these mice (Fig 2C). Unlike *ATOH1*, the gene expression of another intestinal secretory lineage transcription factor, *Hnf1α*, was not significantly changed in CTE mice compared with controls (Fig 2B). ATOH1 downstream signaling was also evaluated at the mRNA level, where we observed a significant decrease in Krüppel-like factor 4 (*KLF4*) expression in CTE mutant mice while the expression of Neurogenin 3 (*NEUG3*) and Sox9 were unchanged compared to controls. We also evaluated factors that promote absorptive lineage differentiation. Although there were no changes in expression of mRNA for Notch (Fig 2B) and its downstream transcription factor Hes1 in CTE mice (Fig 2B & Fig 2C), increased protein expression of notch intracellular domain (NICD) (Fig 2C), the active signal transducer of absorptive lineage specification, corresponded with a bias towards absorptive differentiation. In summary, cell differentiation signaling studies in CTE mice indicate a bias towards absorptive and away from secretory lineage IEC differentiation. This is consistent with the increased number of non-secretory epithelial cells and decreased secretory cells in CTE mice as described [8] and expanded here (Fig 1D). The current study also accords with our previous work in CTE enteroids that showed reduced secretory differentiation [8].

### Enterocytes lack expression of key functional markers

Although our cell differentiation results suggested an increase in absorptive lineage cells in CTE mice, the pathological features seen in CTE patients indicate defective absorption. The secretory nature of diarrhea in CTE patients led us to study ion transporters characteristic of absorptive enterocytes. Moreover, our findings of malabsorption and lack of brush border enzymes responsible for carbohydrate digestion in CTE mice also suggested malfunction of absorptive enterocytes despite an increase in their number. We hypothesized that although enterocytes are increased in number in CTE mice, they are not fully functional due to a lack of key proteins. Hence, additional molecular and functional markers of typical absorptive enterocytes were assessed in the CTE mice. mRNA studies for apical markers showed a decrease in the ion transporter Downregulated in Adenoma (*DRA)* and the water channel aquaporin 7 (*AQP7)* in CTE mutant mice, while mRNA for another apical transporter, sodium-hydrogen exchanger 3 (*NHE3)* and the enterocyte structural marker, villin, were unchanged compared with control mice (Fig 3A). Evaluating basolateral transporters, we found a significant reduction in mRNA levels for anion exchanger 2 (*AE2*) and both alpha and beta subunits of Na, K, ATPase (ATPA1, *ATPB1*) in CTE murine mice while the expression of sodium bicarbonate cotransporter 1 (*NBC1*) was unchanged compared with controls (Fig 3B). These data suggest altered expression of some, but not all, ion transporters in mutant EpCAM enterocytes, and could explain defective water and electrolyte absorption in this disease. The decreased ability of mutant mice to absorb glucose, as shown by the OGTT results, also led us to examine expression of glucose transporters in the small intestine. mRNA levels for sodium-glucose transporter 1 (*SGLT1*) and glucose transporter 2 (*GLUT2*), which are responsible for the majority of glucose transport from the small intestinal lumen to blood, were significantly decreased in CTE murine intestine compared to controls. Additionally, expression of glucose transporter 5 (*GLUT5),* a major fructose transporter of the small intestine, was also significantly decreased in the CTE murine intestine compared with controls (Fig 3D). On the other hand, the lack of change in mRNA expression for sodium-glucose transporter 2 (*SGLT2*) and glucose transporter 1 (*GLUT1*) as well as for villin between the two groups of mice implied the presence of equivalent brush border membranes between the groups. Altered expression of glucose transporters might explain the compromised glucose absorption seen in CTE mutant mice. Decreased disaccharidase activity in CTE mice also prompted investigation of expression of enterocyte-specific genes that aid in digestion and maintaining physiological homeostasis. Expression of enzymes involved in carbohydrate digestion, including maltase-glucoamylase (*MGAM*), lactase (*LAC*), and sucrase-isomaltase (*SI*), was found to be significantly decreased in CTE mice compared to controls (Fig 3C). On the other hand, expression of other villous epithelial markers, such as intestinal alkaline phosphatase (Alk-Phos) and fatty acid binding protein 2 (Fabp2) was equivalent between the two groups. These gene expression results support the findings of selectively decreased disaccharidase activity in CTE mice. Therefore, enterocytes in CTE lack key functional markers, which may contribute to disease pathology.

**Figure 3.**
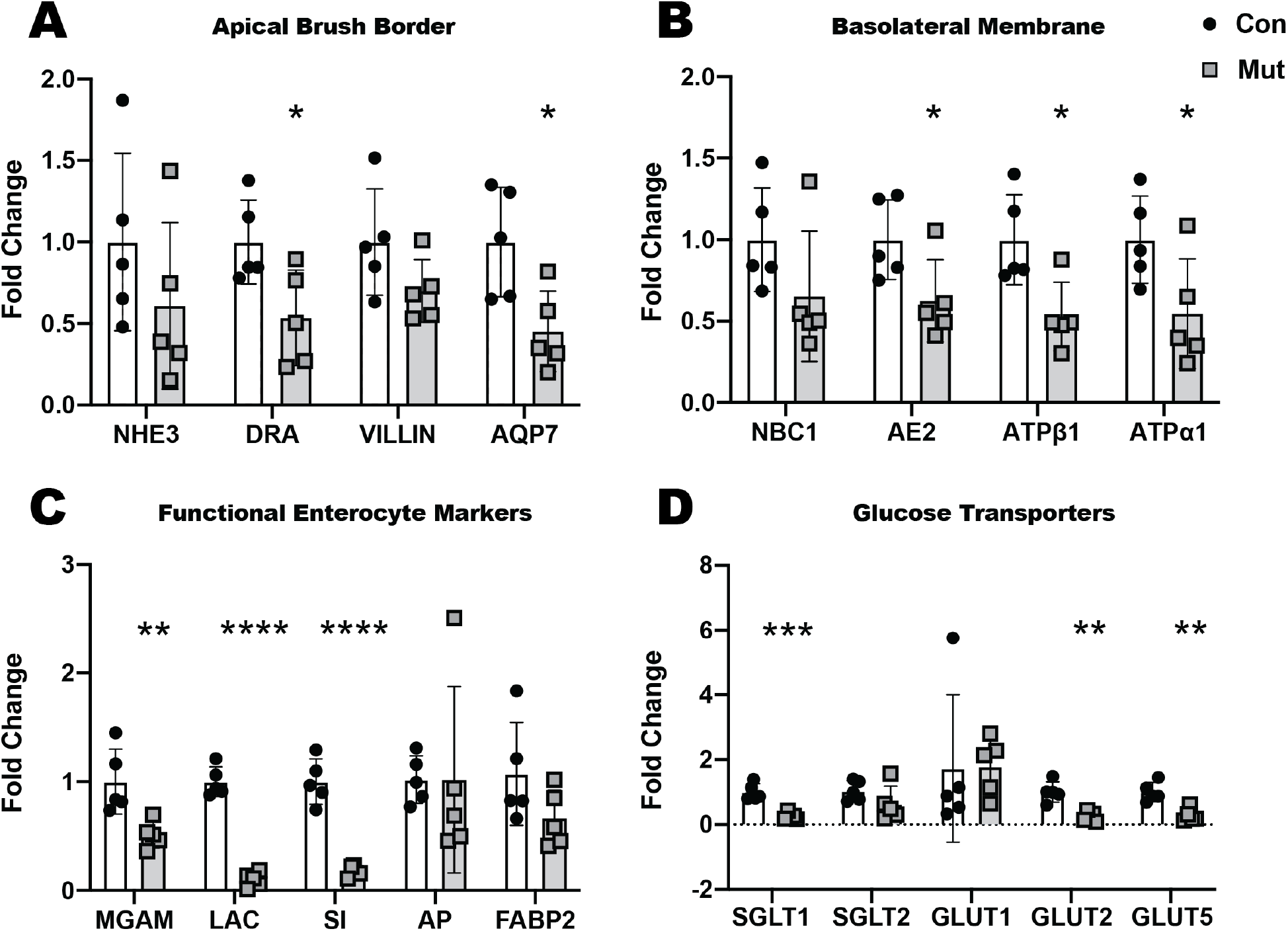
Functional enterocyte gene markers in CTE mice compared with controls. **(A)** Decreased mRNA expression for apical brush border markers Downregulated in Adenoma (*DRA)* and Aquaporin *7 (AQP7)* with normal levels of Sodium hydrogen exchanger 3 (*NHE3*) *and VILLIN* in intestinal tissue lysates of mutant (Mut) mice compared to control (Con) mice (n=5). **(B)** Decreased mRNA expression for basolateral Anion exchanger 2 *(AE2),* ATP-ase beta 1 (*ATPβ1)* and ATP-ase alpha 1 (*ATPα1*) in intestinal tissue lysates of Mut when compared to Con mice (n=5) with normal levels of membrane marker Sodium bicarbonate cotransporter 1 (*NBC1).* **(C)** Decreased mRNA expression for functional enterocyte markers Maltase glucoamylase (*MGAM),* Lactase *(LAC),* Sucrase Isoamylase (*SI*) with normal expression of Alkaline phosphatase *(AP)* and Fatty acid binding protein *2 (FABP2)* in intestinal tissue lysates in Mut mice compared to Mut mice (n=5). **(D)** Decreased mRNA expression of glucose transporters Sodium glucose cotransporter 1 *(SGLT1),* Glucose transporters 2 and 5 (*GLUT2, GLUT5*) and normal expression of *SGLT2* and *GLUT1* in intestinal tissue lysates in Mut compared to Con mice (n=5). *=p<0.05, **=p<0.01, ***=p<0.001, ****=p<0.0001 by Student’s t test for all results.

Due to evidence for altered gene expression of various apical and basolateral membrane markers, including glucose transporters, in Mut mice, protein expression was evaluated by western blot. The structural protein, villin, and the ion transporter, DRA, were present at comparable levels in both control and mutant mice. Protein expression for Villin is in line with its mRNA expression that confirms the expression of villin is unchanged between control and CTE mutant mice. Although the mRNA expression of DRA is decreased between control and mutant mice, protein expression was found to be equivalent between the groups. This may be due less dynamic DRA protein alteration than mRNA. As both RNA and protein studies were performed at a single time point, it is possible that sustained decrease of DRA mRNA has yet to downregulate the DRA protein expression. Importantly, protein levels of NHE3, and SGLT1 were significantly decreased in mutant mice compared to controls (Fig 4A). These results further confirm the deficiency of key markers in CTE-affected enterocytes.

**Figure 4.**
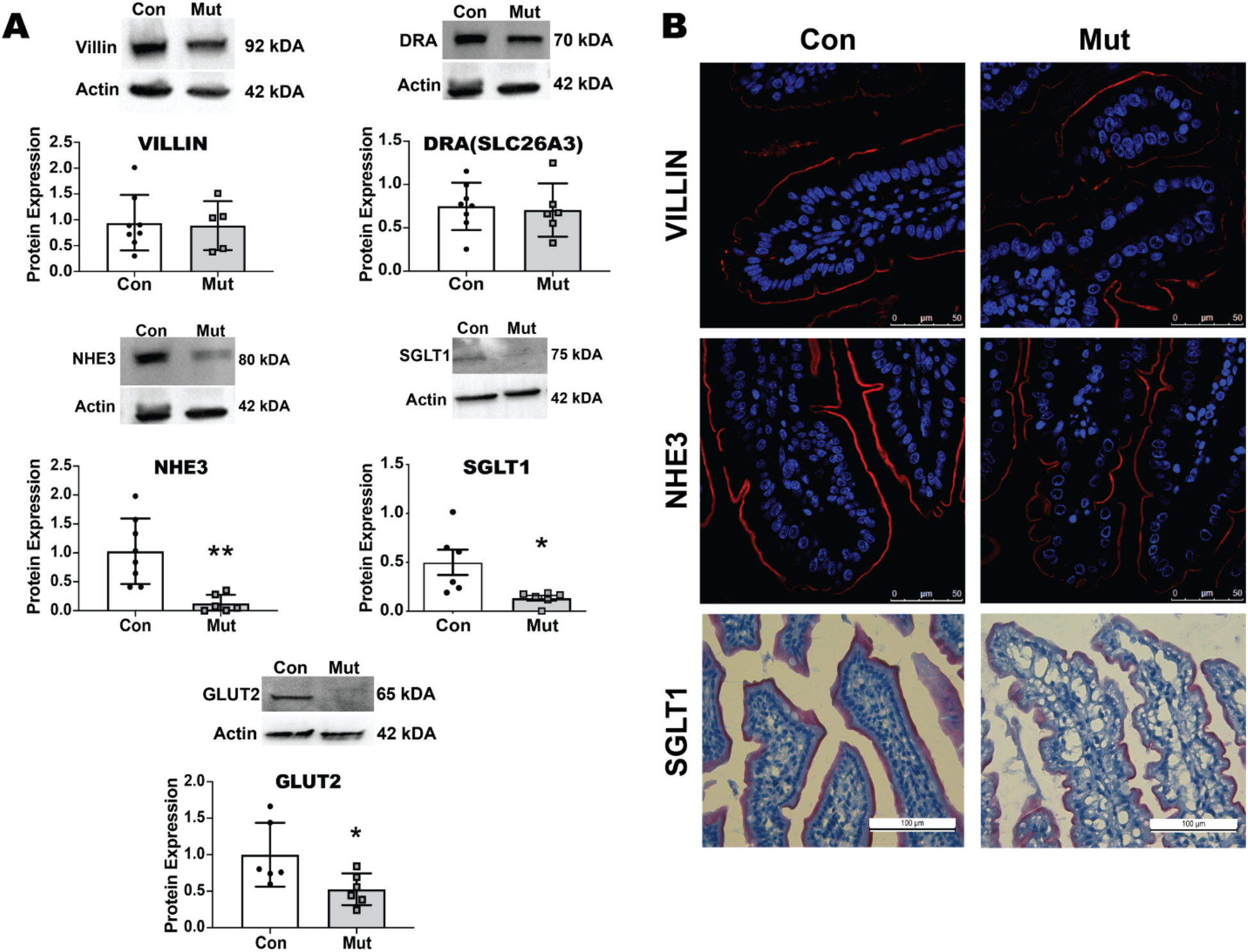
Functional enterocyte markers at protein level in CTE mice. **(A)** Representative western blot images and quantification of VILLIN, Downregulated in Adenoma (DRA), Sodium hydrogen exchanger 3 (NHE3), Sodium glucose co transporter 1 (SGLT1) and Glucose transporter 2 (GLUT2) with respect to β-actin (Actin) in intestinal tissue lysates from Control (Con) and CTE mutant (Mut) mice, with decreased in NHE3, SGLT1 and GLUT2 expression and unchanged VILLIN and DRA expression. The same blot was used to detect DRA and NHE3 protein. *=p<0.05, **=p<0.01, ***=p<0.001, ****=p<0.0001 by Student’s t test for all results. **(B)** Representative images of immunofluorescent staining of Con and Mut intestinal tissue section showing expression of VILLIN (Red) and Draq5 staining for nuclear DNA (Blue) (upper panel); expression of NHE3 (Red) and Draq5 staining for nuclear DNA (Blue) (middle panel). Representative brightfield images of immunohistochemical staining of Con and Mut intestinal tissue sections showing expression of SGLT1. NHE3 and SGLT1 lack uniform distribution along the brush border membrane in Mut tissue when compared to Con while there was no change in localization of VILLIN expression along the membrane (n=3). Scale bar for all immunofluorescence images represents 50 μm; Scalebar for brightfield images denotes 100 μm.

The decrease in gene and protein expression and disruption in the function of absorptive enterocytes led us to question the location of enterocyte membrane proteins in the intestinal epithelium. In order to test this, immunofluorescent (IF) and IHC staining was performed for NHE3, villin, and SGLT1 on small intestinal sections from control and mutant CTE mice. No changes were found in the expression and localization of villin between the groups, confirming an intact brush border. Conversely, staining of SGLT1 and NHE3 was abberant in CTE mice (Fig 4B). Both markers lacked uniform staining along the brush border membrane in CTE mice, whereas uniform expression of these markers was noted throughout the brush border membrane in control mice.

### Notch pathway inhibition increased secretory cell markers in CTE enteroids

Our findings of aberrant epithelial differentiation in CTE mice, resulting in an increase in the absorptive lineage and decreased secretory lineages as well as the identification of potentially affected signaling pathways led us to investigate whether these discrepancies could be rescued by targeting relevant signaling pathways in CTE mice. Because Notch intercellular domain was increased in CTE epithelial cells (Fig 2C), Notch signaling was targeted. Notch pathway signaling was inhibited with the γ-secretase inhibitor, DAPT, in CTE enteroids. This resulted in an increase in mRNA for the goblet cell marker, MUC2, and the enteroendocrine cell marker, ChgA, compared to mutant enteroids without any treatment; however, mRNA for the Paneth cell marker, lysozyme, was not significantly altered (Fig 5A). Expression of transcription factors relevant to the secretory and absorptive pathway was also examined. *ATOH1* and *Neug3* were increased in DAPT-treated mutant enteroids compared to untreated Mut enteroids, while Hes1 expression was not affected. These results suggest that aberrant Notch signaling may be targeted to improve the balance of cell differentiation in CTE.

**Figure 5.**
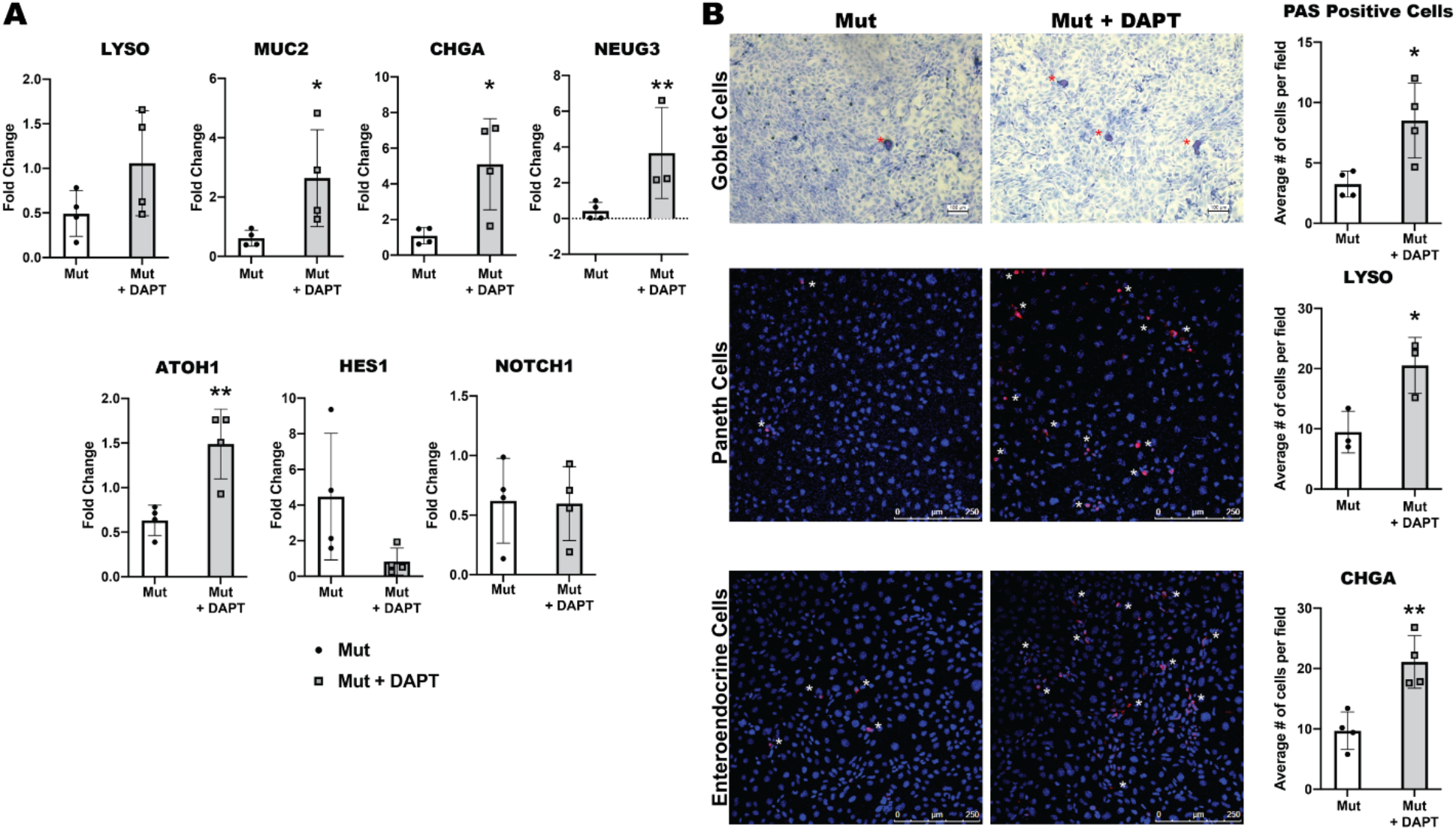
Restoration of secretory epithelial lineages with Notch inhibition in CTE enteroids. **(A)** mRNA expression for cell type markers (*LYSO, MUC2, CHGA, NEUG3*) and transcription factors (*ATOH1, HES1, and NOTCH*) in lysates from mutant enteroids (Mut) with or without treatment with DAPT (n=4). The expression of *MUC2, CHGA, NEUG3*, and *ATOH1* improved upon treatment with DAPT, while expression of *LYS, HES1* and *NOTCH* was not changed. **(B)** Representative bright field of PAS staining for goblet cells (n=4) and confocal images of Lysozyme for Paneth Cells (n=3) and Chromogranin A for enteroendocrine cells (n=4) with quantification of cell types. Red/White asterisks denote the positive cells in the respective images. Increased number of goblet, Paneth and enteroendocrine cells were found in DAPT treated Mut enteroids when compared with untreated Mut enteroids. For bright field image, scale bar denotes 100μm; for immunofluorescent images, the scale bar represents 250μm. *=p<0.05, **=p<0.01, ***=p<0.001, ****=p<0.0001 by Student’s t test for all results.

To confirm these results, markers of secretory cells were evaluated at the protein level. In line with the mRNA findings, DAPT increased the number of secretory cells in mutant enteroid-derived monolayers (EDM). Thus, the proportions of lysozyme-positive (Paneth cells), PAS-positive (goblet cells) and chromogranin A-positive (endoenterocrine) cells were significantly increased in DAPT-treated mutant EDMs compared to untreated mutant EDMs (Fig 5B). This result reveals a potential therapeutic target to restore secretory cells in CTE disease.

### CTE-associated changes in epithelial homeostasis primarily affected the small intestine

Although, CTE manifests primarily in the small intestine of human patients and results in severe intestinal failure [2], the colonic mucosa has also been reported to be involved [1]. We therefore examined whether EpCAM mutation impacts epithelial biology in two other GI epithelial organs, the colon and pancreas. To evaluate the effect of CTE in the colon, hematoxylin and eosin (H&E) staining as well as qRT-PCR of glucose transporters, and markers of the apical brush border were performed. H&E staining revealed that the overall architecture of the colon was preserved in mutant mice compared to controls. The colonic epithelium displayed mild cellular disorganization along the brush border in mutant mice compared to controls. No change was found in mRNA expression for any of the markers studied between control and mutant mice (Supplemental Fig S1). In the pancreas, H&E staining revealed no change in acinar cells, islets of Langerhans, or interlobular ducts between control and mutant mice (Supplemental Fig S2). Finally, to further investigate whether EpCAM mutants influence pancreatic activity, serum lipase and amylase were evaluated in control and mutant mice. Equivalent serum lipase and amylase activity in control and mutant mice (Supplemental Fig 2) suggests pancreatic function is likely to be normal in CTE mice. These experiments confirm that CTE most severely targets the small intestine.

## Discussion

Previously, studies in mutant EpCAM murine models have revealed alterations in histopathology, epithelial barrier and ion transport function, and decreased secretory cells that contribute to CTE pathology [6, 7]. Studies have also suggested decreased secretory epithelial cells in different CTE models (patient, mice, enteroids) [8], though alterations in the differentiation cascade that leads to differentially expressed cell types had not been investigated. Moreover, despite knowledge of the classical pathological features in CTE, their relationship to the nutrient malabsorption observed in patients [1] had not been studied in any CTE model. The biology behind the signs and symptoms in CTE patients, such as impaired growth and the secretory, as well as osmotic nature, of diarrhea, remained elusive. The current study shows that CTE murine model, bearing a mutation in EpCAM, exhibits alterations in small intestinal epithelial diffentiation. The aberrant lineage differentiation cascade in CTE mice leads to decreased secretory and increased absorptive epithelial cells. Moreover, treatment of our enteroid model with the γ-secretase inhibitor, DAPT, provides a therapeutic suggestion to rescue faulty lineage differentiation. Our study also suggests that dysfunctional absorptive enterocytes are a major contributor to pathology characteristic of CTE, such as malabsorption, reduced caloric uptake, secretory and osmotic diarrhea, and weight loss.

Failure to thrive and impaired growth are hallmarks of CTE [1]. A lack of weight gain has been reported in constitutive CTE neonatal mice [6]. In the current study, weight loss also validates the adult model system. Failure to thrive and weight loss suggest undernutrition due to inadequate caloric intake, absorption, and/or excessive energy expenditure. The growth failure in CTE mice provided an impetus for an evaluation of malabsorption. The clinical observation that enteral feeding often worsens the symptoms of CTE and parenteral nutrition being the most reliable source of nutrients in these patients, also indicates the involvement of malabsorption in the disease [1]. Since impaired growth worsens disease outcome because it leads to complications, treatments that alleviate weight loss may improve prognosis.

Lack of glycoside hydrolases (maltase, sucrase and lactase) in the small intestinal brush border in CTE mice might lead to reduced digestion and availability of glucose for absorption, resulting in malabsorption of ingested calories. Since CTE appears in the neonatal period when milk is the major source of nutrient intake, a significant decrease in lactase activity in our CTE mice is consistent with impaired growth in CTE patients and models of the disease [1, 33, 34]. Moreover, both the lack of digestive enzyme activity and glucose malabsorption likely contribute to the osmotic diarrhea reported in CTE patients [35]. Although beyond the scope of this study, evaluation of additional hydrolases synthesized and secreted by the pancreas into the luminal content would provide further evidence as to whether brush border enzymes are selectively affected in CTE. Nevertheless, since pancreatic structure and function are not affected in CTE, it is likely that pancreatic secreted enzymes would be normal in CTE mice, consistent with our finding of normal levels of circulating lipase and amylase. On the other hand, the increase of a ubiquitous intestinal enzyme activity, alkaline phosphatase, in the CTE mice is interesting as it suggests that the intestinal epithelium is still largely intact despite a decrease in one of its major functions.

Enteroendocrine cells comprise the largest endocrine system in the body and regulate physiological and homeostatic function of the digestive tract by secreting multiple peptide hormones [36]. Thus, a decrease in enteroendocrine cell number in CTE mice might disrupt sensing of luminal content and activation of vagal afferent fibers to control food intake [37]. A lack of intestinal enteroendocrine cells is not unique to CTE, and is also seen in congenital anendocrinosis [38]. As with CTE, patients with congenital anendocrinosis have severe nutrient malabsorption leading to diarrhea.

ATOH1 is the master regulator that directs progenitor cells to the secretory cell fate [39]. Hence, the novel finding of decreased ATOH1 stemming from EpCAM mutation likely accounts for the aberrant cell census found in CTE [8]. On the other hand, an increase in NICD in CTE mice confirms that IEC differentiation in CTE drives differentiation away from secretory cell fate towards the absorptive lineage. Rescue of the decreased secretory cells with DAPT treatment further confirms involvement of the Notch pathway in CTE and also potentially opens an avenue towards future targeted therapy in CTE. Further research to explore this option is needed.

The increased Notch signaling in adult CTE mice explains the increased number of non-secretory epithelial cells and upregulated alkaline phosphatase activity found in these mice. Of note, an increase in epithelial proliferation was reported in the neonatal CTE model [6]. The increased number of absorptive epithelial cells in adult CTE mice along with unchanged expression of Cdx2 and several other epithelial markers not only excludes a general detrimental effect of CTE on all IEC types but also strongly supports an alteration in the IEC differentiation cascade. Since EpCAM is involved in pluripotency and maintenance of ISCs, the decreased expression of Lgr5 in CTE mice is probably due to a direct effect of EpCAM mutation. ER stress is also reported to be responsible for loss of intestinal epithelial stemness [40] through activation of the unfolded protein response (UPR). Our prior finding of ER stress and UPR in CTE mice [41] may be the cause of of the loss of Lgr5-positive ISCs in these mice. Further studies are needed to explain the direct mechanism by which mutant EpCAM affects Lgr5-positive ISCs but not other stem cells, transient amplifying cells or other progenitor cells.

Our findings suggest some key markers of mature absorptive enterocytes are inappropriately expressed in CTE mice while expression of other markers is not changed. Decreased expression of key ion transporters/channel/pumps at both basolateral and apical membranes may account for secretory diarrhea found in CTE patients. Reduced expression of SGLT1 and GLUT2 would lead to glucose malabsorption and less caloric intake in these CTE mice. Though it is beyond the scope of current study to fully explain aberrant expression/localization of some but not all markers in affected enterocytes, we can speculate upon some possible mechanisms. For example, EpCAM contributes to the formation of a functional apical junctional complex with adaptor molecules that play a role in regulating cell polarity [42]. Thus, the observation that NHE3 protein, but not at mRNA, expression may imply post transcriptional regulation via altered interactions with the PDZ domain protein, NHERF [43]. Moreover, we recently reported that endoplasmic reticulum stress and the unfolded protein response (UPR) play a role in pathogenesis in mutant EPCAM mice [41]. Activation of UPR might lead to alterations in polarized absorptive enterocytes, which normally depend on intracellular transport of various components of the brush border for proper functioning.

The current study enhances our understanding of the pathogenesis of CTE and the role of mutant EpCAM in the disease. There are currently no cures nor directly-targeted therapies for patients with this devastating disease, and thus understanding disease mechanisms is a necessary step towards improving disease outcomes. Future studies may also reveal how the cascade of mechanisms that regulate intestinal epithelial differentiation intersect with mutant EpCAM signaling pathways in CTE. Targeting of the biological processes uncovered here will lay the groundwork for the expansion of CTE therapeutics.

## Materials and Methods

### Animals

All animal experiments were performed according to the instructions and guidelines of, and were approved by, the University of California San Diego (UCSD) Institutional Animal Care and Use Committee (IACUC). Control C57BL/6N mice were obtained from Charles River (Wilmington, MA). A previously described adult murine model with tamoxifen-inducible expression of mutant EpCAM was used for the current study (Mut) [7]. Mutant EpCAM was expressed upon oral gavage of tamoxifen. Over the course of 5 days, mice were given tamoxifen on days 1 and 2 and had their weights measured daily. Studies were primarily performed at day 5. In addition, littermates with wild type EpCAM were orally gavaged with tamoxifen alongside the Mut mice to serve as controls (Con) for all experiments. Mice were studied between 9-14 weeks of age. All mice were euthanized using carbon dioxide (CO_2_) gas asphyxiation in a CO_2_ chamber followed by cervical dislocation. Both male and female mice were used in the study to avoid sex bias.

### Enteroid Culture

Enteroid cultures were initiated, maintained, and expanded as previously described [8]. For EDMs, enteroids were disrupted and plated on 6.5-mm Transwell inserts with 0.4-μm pores (Corning Inc, Corning, NY) with 5% conditioned media (CM) for 2 days as described [8]. 3 μM tamoxifen was added to the 5% CM media to generate CTE mutant EDM. For inhibition of the Notch signaling pathway, 100 μM N-[N-(3,5-Difluorophenacetyl)-L-alanyl]-S-phenylglycine t-butyl ester (DAPT) (AdipoGen Life Science, San Diego, CA) was added to the culture media simultaneously with tamoxifen treatment.

### Oral Glucose Tolerance Test (OGTT)

All mice were fasted for 4 hours with free access to drinking water prior to being challenged with 2g/kg of glucose by oral gavage, using 30% glucose prepared in drinking water. Blood was sampled via the tail vein at 0, 10, 30, and 90 minutes post glucose administration. Glucose concentration in blood was measured using an glucometer (Freestyle Lite, Abbott, Chicago, IL) and expressed as mg/dL.

### Enzyme Activity Assay

Frozen duodenal tissue with luminal content was homogenized as follows. After weighing, the tissue was crushed with a plastic pestle in a nine-fold volume of PBS supplemented with protease inhibitors in a tube kept on ice. Following centrifugation at 3500xg for 10 min at 4°C, the supernatant was used for various assays and the protein content of the homogenate was measured using DC protein assay kit (Bio-Rad, Hercules, CA). Lactase activity was assessed using a Lactase Assay Kit (Cat# E-BC-K131-S, Elabscience, Houston, TX) according to the manufacturer’s instruction. Alkaline phosphatase activity was assessed using an alkaline phosphatase fluorometric assay kit (ab83371, Abcam, Cambridge, UK) according to manufacturer’s instruction. Maltase and sucrase activity were measured colorimetrically using the glucose oxidase peroxidase method with some modification of a published protocol [44]. Briefly, 10 μl of the tissue homogenate were added to 10 μl of the substrate (56 mM sucrose or maltose) and incubated for 1 hour at @37°C for generation of glucose. The quantity of glucose produced was then measured using a Glucose assay Kit (Sigma-Aldrich, St. Louis, MO; #GAGO20). For all assays, enzyme activity was normalized to total protein content.

### Paraffin and Frozen Section Preparation

Jejunal sections were fixed in 10% neutral buffered formalin (Fisher Scientific, Waltham, MA) containing zinc (for paraffin) or 4% paraformaldehyde (for frozen sections) overnight at 4°C and embedded in paraffin or Tissue-Tek Optimal Cutting Temperature (OCT) (Sakura Finetek USA, Torrance, CA), respectively. Tissue for frozen sectioning was cryoprotected in a 30% sucrose solution (Macron, Center Valley, PA) for an additional night at 4°C. 10 μm sections were utilized.

### Immunofluorescent (IF) staining

Immunofluorescent staining was performed in both intestinal tissue sections and EDMs. To stain intestinal sections, paraffin-embedded sections were deparaffinized and hydrated prior to blocking steps. Antigen retrieval was performed at pH 7 via microwave (VECTOR Antigen Unmasking Solution, Vector labs, Burlingame, CA) before further processing. EDMs were cultured in chamber slides in a fashion to similar to that used for transwell inserts. Cell monolayers were fixed with 4% PFA for 3 min. Both intestinal tissue and EDM samples were blocked in 5% goat serum in tris-buffered saline (TBS) (tissue staining) or in phosphate buffered saline (PBS) (EDM) for 1 hour at room temperature. Sections were incubated overnight with primary antibodies at 4°C in blocking buffer. Samples were then washed thrice for 5 minutes in TBS (tissue)/PBS (EDM) and incubated in Alexa Fluor 568 goat anti-rabbit antibodies (Thermo Fischer Scientific, Waltham, MA) for 1 hour at room temperature. Nuclei were stained using the Draq5 fluorescent probe (Thermo Fischer Scientific) in TBS (tissue staining)/PBS (EDM) and incubated for 10 minutes at room temperature. Finally, the samples were washed in PBS and mounted using Prolong Gold antifade reagent (Life Technologies, Eugene, OR). Slides were imaged using a Leica (Leica Microsystems, Buffalo Grove, IL) confocal imaging system (DMI4000 B) using a 25× Plan-Apo 0.8 numerical aperture with 40X objective lens. Images were captured using LASX v4.1 Image Acquisition software supplied by Leica.

The primary antibodies used were as follows: NHE3 (GTX41967, GeneTex, Irvine, CA), Villin-1 (2369S, Cell Signaling Technology, Danvers, MA), Lysozyme (PA5-16668, Thermo Fisher Scientific), Chromogranin A (ab15160, Abcam).

### Immunohistochemical (IHC) staining

For immunohistochemical analysis of paraffin-embedded sections, endogenous peroxidase activity was blocked with a 3% H_2_O_2_ solution (VECTOR Antigen Unmasking Solution Citrate-based pH 6.0 H-3300, Vector labs) prior to blocking with 5% goat serum in TBS for 1 hour at room temperature. The sections were then incubated with primary antibodies in blocking buffer overnight at 4°C. Samples were washed thrice for 5 minutes in TBS. After washing, slides were incubated with secondary antibodies for 30 minutes at room temperature. The signal was revealed using Vectastain Elite ABC-HRP kit and AEC Peroxidase (HRP) substrate 3-amino-9ethylcarbazole (VECTOR AEC Peroxidase Subtract Kit SK-4200, Vector Laboratories, Burlingame, CA). Samples were mounted using ProLong Gold antifade reagent (Life Technologies, Eugene, OR). Slides were imaged using a Leica (Leica Microsystems, Buffalo Grove, IL) inverted imaging system (DMi1) using a Hi Plan I 0.30 and 0.50 numerical aperture with 20X objective lens. Images were acquired at 20x magnification for enteroendocrine cell counts. Two representative images were taken of each section. All positively stained cells were counted within the image, followed by a count of all epithelial cells present within the image. The number of positively stained cells was then divided by total epithelial cells to generate a percentage of positive cells for one image, which was then averaged with the second image’s percentage to create an average for each mouse.

Frozen sections were used for the immunohistochemical analysis of SGLT1. Slides were blocked in 5% rabbit serum and 1.5% BSA in TBS containing 0.2% Triton X-100 (TBST). The primary antibody was diluted in TBST overnight at 4°C. Samples were washed for 10 minutes 3 times in TBS containing 0.0005% Triton X-100. Slides were incubated in goat secondary for 30 minutes at room temperature. The signal was revealed using Vectastain Elite ABC-AP kit and Alkaline Phosphatase (AP) substrate (VECTOR Red Alkaline Phosphatase Substrate Kit SK-5100, Vector Laboratories). Samples were mounted using Prolong Gold antifade reagent (Life Technologies, Eugene, OR). Slides were imaged using Leica (Leica Microsystems, Buffalo Grove, IL) inverted imaging system (DMi1) using a Hi Plan I 0.30 and 0.50 Numerical aperture with 20X objective lens. The primary antibodies used were as follows: Chromogranin A (ab15160, Abcam), SGLT1 (EB09310, Everest Biotech, Oxfordshire, UK).

### Periodic Acid-Schiff Staining

Periodic acid-Schiff (PAS) staining was performed to stain mucin-producing goblet cells in murine EDMs using a PAS Kit (Sigma-Aldrich) as described previously [8]. Briefly, EDMs in chamber slides were fixed with 4% PFA for 3 minutes and stained with periodic acid for 1 minute, followed by Schiff’s reagent for 5 minutes. The monolayer was then counterstained with hematoxylin for 1 minute. Scott’s bluing reagent was used after hematoxylin staining in both cases. Cells that stained magenta were considered PAS positive for goblet cell counts.

### Western Blotting

Intestinal tissue extracts were prepared by cell lysis using RIPA buffer (CST), followed by centrifugation (14,000 g for 15 minutes) at 4°C. Proteins were resolved on a 4-12% SDS-PAGE gel, followed by transfer to a PVDF membrane (Immobilon-PSQ PVDF membrane, Millipore-Sigma; Burlington, MA). Immunoblotting was performed as described earlier [41], using rabbit polyclonal antibodies to HES1 (Thermo Fisher Scientific), MATH1 (ATOH1) (Developmental Studies Hybridoma Bank), NICD (Abcam, Cambridge, UK), NHE3 (GTX41967, GeneTex), SGLT1 (EB09310 Everest Biotech) and GLUT2 (ab54460, Abcam). Mouse anti-β-Actin staining (A1978; Sigma-Aldrich) served as a loading control. The band intensities from the western blot images were analyzed with Image J after normalizing for the loading control and the fold change of protein expression in mutant models was calculated relative to the control group.

### Real-Time qRT-PCR

Total mRNA for Real-Time quantitative (q)RT-PCR studies of small intestine of control and mutant mice was isolated as per the manufacturer’s instruction using a quick RNA micro prep kit (Zymo research) as described previously [8]. First-strand cDNA was synthesized from 250 ng of total RNA with iScript cDNA Synthesis kit (Bio-Rad) following the manufacturer’s protocol. The primers used for Real-Time qRT-PCR in the current study are listed in Table 1. Real-time qRT-PCR reactions were set up using FastStart Universal SYBR Green Master Mix (Thermo Fisher Scientific) and thermal cycling was performed using a StepOnePlus (Applied Biosystems) Real-Time PCR System using Step One software v2.0 (Applied Biosystems). All qRT-PCR reactions were performed in duplicate. The relative fold change of the respective gene was calculated after normalization to the housekeeping gene (18s rRNA) and comparison to the control group.

**Table 1.**
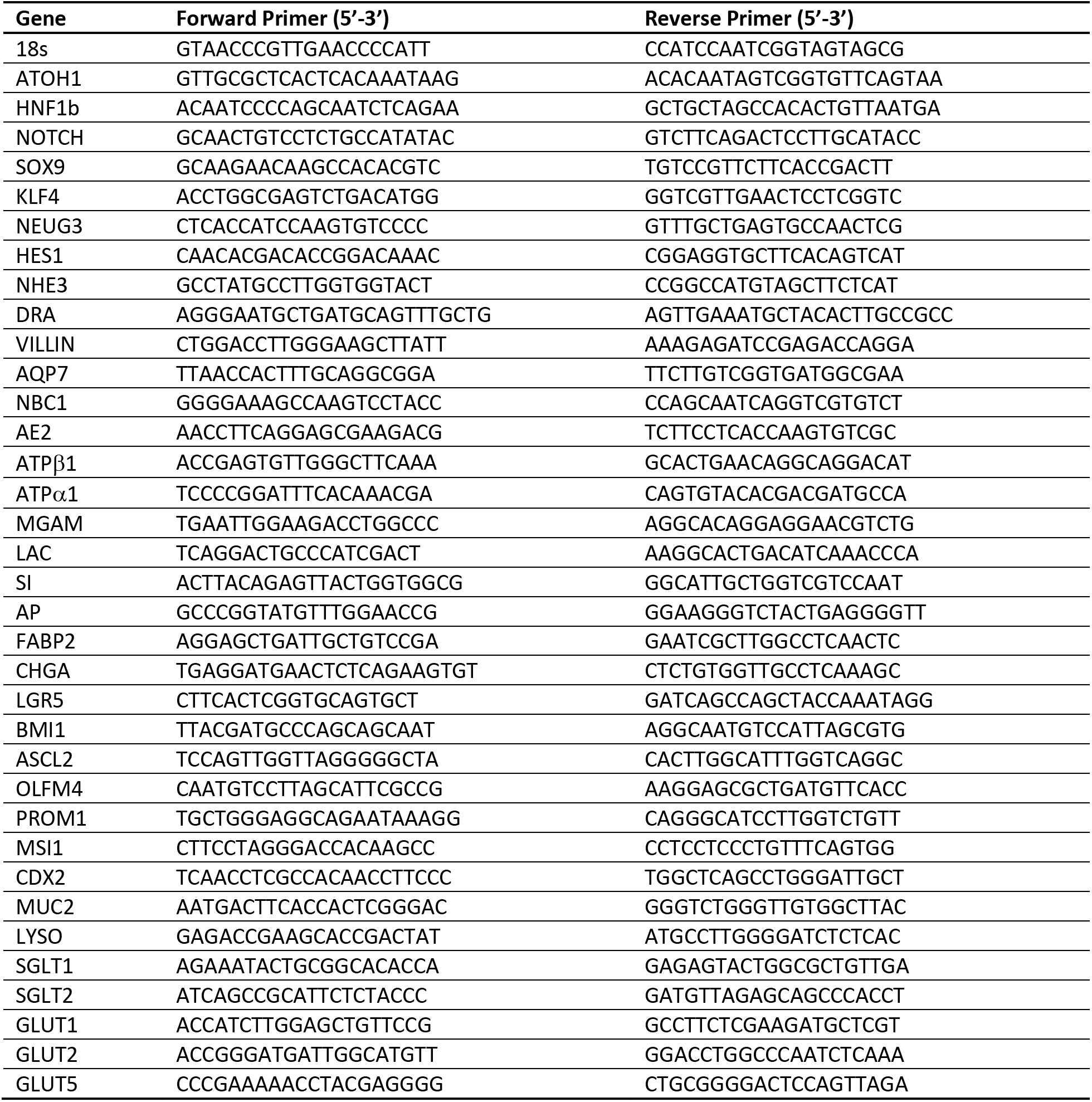
Primer sequences.

### Statistical Analysis

Data are presented for at least three biological replicates unless otherwise indicated. Error bars represent the standard deviation (SD) of the mean of the measured parameter. The statistical significance of differences between the two groups was calculated by Student’s unpaired t-test (Prism 7; GraphPad software Inc). Two-tailed P values of <0.05 were considered statistically significant. P values were designated as P>0.05; *, P < 0.05; **, P < 0.01 and ***, P < 0.001.

## Supporting information

Supplemental results

## Acknowledgments

We would like to thank Hal Hoffman, Lawrence Prince, Ben Croker, Soumita Das, and Lori Broderick for useful discussion. The graphical abstract figure was created with BioRender.com.

## References

[1] Goulet O, Salomon J, Ruemmele F, de Serres NP-M, Brousse N. Intestinal epithelial dysplasia (tufting enteropathy). Orphanet journal of rare diseases 2007;2(1):20.

[2] Reifen RM, Cutz E, Griffiths A-M, Ngan BY, Sherman PM. Tufting enteropathy: a newly recognized clinicopathological entity associated with refractory diarrhea in infants. Journal of pediatric gastroenterology and nutrition 1994;18(3):379–85.

[3] Sivagnanam M, Mueller JL, Lee H, Chen Z, Nelson SF, Turner D, Zlotkin SH, Pencharz PB, Ngan BY, Libiger O, Schork NJ, Lavine JE, Taylor S, Newbury RO, Kolodner RD, Hoffman HM. Identification of EpCAM as the gene for congenital tufting enteropathy. Gastroenterology 2008;135(2):429–37.

[4] Pathak SJ, Mueller JL, Okamoto K, Das B, Hertecant J, Greenhalgh L, Cole T, Pinsk V, Yerushalmi B, Gurkan OE, Yourshaw M, Hernandez E, Oesterreicher S, Naik S, Sanderson IR, Axelsson I, Agardh D, Boland CR, Martin MG, Putnam CD, Sivagnanam M. EPCAM mutation update: Variants associated with congenital tufting enteropathy and Lynch syndrome. Hum Mutat 2019;40(2):142–61.

[5] Guerra E, Lattanzio R, La Sorda R, Dini F, Tiboni GM, Piantelli M, Alberti S. mTrop1/Epcam knockout mice develop congenital tufting enteropathy through dysregulation of intestinal E-cadherin/β-catenin. PLoS One 2012;7(11):e49302.

[6] Mueller JL, McGeough MD, Pena CA, Sivagnanam M. Functional consequences of EpCam mutation in mice and men. Am J Physiol Gastrointest Liver Physiol 2014;306(4):G278–88.

[7] Kozan PA, McGeough MD, Peña CA, Mueller JL, Barrett KE, Marchelletta RR, Sivagnanam M. Mutation of EpCAM leads to intestinal barrier and ion transport dysfunction. Journal of Molecular Medicine 2015;93(5):535–45.

[8] Das B, Okamoto K, Rabalais J, Kozan PA, Marchelletta RR, McGeough MD, Durali N, Go M, Barrett KE, Das S, Sivagnanam M. Enteroids expressing a disease-associated mutant of EpCAM are a model for congenital tufting enteropathy. Am J Physiol Gastrointest Liver Physiol 2019;317(5):G580–G91.

[9] Barth AIM, Kim H, Riedel-Kruse IH. Regulation of epithelial migration by epithelial cell adhesion molecule requires its Claudin-7 interaction domain. PLoS One 2018;13(10):e0204957.

[10] Salomon J, Gaston C, Magescas J, Duvauchelle B, Canioni D, Sengmanivong L, Mayeux A, Michaux G, Campeotto F, Lemale J, Viala J, Poirier F, Minc N, Schmitz J, Brousse N, Ladoux B, Goulet O, Delacour D. Contractile forces at tricellular contacts modulate epithelial organization and monolayer integrity. Nat Commun 2017;8:13998.

[11] Lei Z, Maeda T, Tamura A, Nakamura T, Yamazaki Y, Shiratori H, Yashiro K, Tsukita S, Hamada H. EpCAM contributes to formation of functional tight junction in the intestinal epithelium by recruiting claudin proteins. Dev Biol 2012;371(2):136–45.

[12] Wu C-J, Feng X, Lu M, Morimura S, Udey MC. Matriptase-mediated cleavage of EpCAM destabilizes claudins and dysregulates intestinal epithelial homeostasis. The Journal of clinical investigation 2017;127(2):623.

[13] Noah TK, Donahue B, Shroyer NF. Intestinal development and differentiation. Exp Cell Res 2011;317(19):2702–10.

[14] Fre S, Huyghe M, Mourikis P, Robine S, Louvard D, Artavanis-Tsakonas S. Notch signals control the fate of immature progenitor cells in the intestine. Nature 2005;435(7044):964–8.

[15] Kim TH, Shivdasani RA. Genetic evidence that intestinal Notch functions vary regionally and operate through a common mechanism of Math1 repression. J Biol Chem 2011;286(13):11427–33.

[16] Shroyer NF, Helmrath MA, Wang VY, Antalffy B, Henning SJ, Zoghbi HY. Intestine-specific ablation of mouse atonal homolog 1 (Math1) reveals a role in cellular homeostasis. Gastroenterology 2007;132(7):2478–88.

[17] Zecchini V, Domaschenz R, Winton D, Jones P. Notch signaling regulates the differentiation of post-mitotic intestinal epithelial cells. Genes Dev 2005;19(14):1686–91.

[18] Canani RB, Terrin G. Recent progress in congenital diarrheal disorders. Curr Gastroenterol Rep 2011;13(3):257–64.

[19] Kuokkanen M, Kokkonen J, Enattah NS, Ylisaukko-Oja T, Komu H, Varilo T, Peltonen L, Savilahti E, Jarvela I. Mutations in the translated region of the lactase gene (LCT) underlie congenital lactase deficiency. Am J Hum Genet 2006;78(2):339–44.

[20] Martin MG, Turk E, Lostao MP, Kerner C, Wright EM. Defects in Na+/glucose cotransporter (SGLT1) trafficking and function cause glucose-galactose malabsorption. Nat Genet 1996;12(2):216–20.

[21] Ritz V, Alfalah M, Zimmer KP, Schmitz J, Jacob R, Naim HY. Congenital sucrase-isomaltase deficiency because of an accumulation of the mutant enzyme in the endoplasmic reticulum. Gastroenterology 2003;125(6):1678–85.

[22] Wedenoja S, Pekansaari E, Hoglund P, Makela S, Holmberg C, Kere J. Update on SLC26A3 mutations in congenital chloride diarrhea. Hum Mutat 2011;32(7):715–22.

[23] Dhekne HS, Hsiao NH, Roelofs P, Kumari M, Slim CL, Rings EH, van Ijzendoorn SC. Myosin Vb and Rab11a regulate phosphorylation of ezrin in enterocytes. J Cell Sci 2014;127(Pt 5):1007–17.

[24] Overeem AW, Posovszky C, Rings EH, Giepmans BN, van IJzendoorn SC. The role of enterocyte defects in the pathogenesis of congenital diarrheal disorders. Disease models & mechanisms 2016;9(1):1–12.

[25] Yan KS, Chia LA, Li X, Ootani A, Su J, Lee JY, Su N, Luo Y, Heilshorn SC, Amieva MR, Sangiorgi E, Capecchi MR, Kuo CJ. The intestinal stem cell markers Bmi1 and Lgr5 identify two functionally distinct populations. Proc Natl Acad Sci U S A 2012;109(2):466–71.

[26] Baker M. Intestinal stem cells: one gene to rule them all. Nature Reports Stem Cells 2009.

[27] van der Flier LG, Haegebarth A, Stange DE, van de Wetering M, Clevers H. OLFM4 is a robust marker for stem cells in human intestine and marks a subset of colorectal cancer cells. Gastroenterology 2009;137(1):15–7.

[28] Snippert HJ, van Es JH, van den Born M, Begthel H, Stange DE, Barker N, Clevers H. Prominin-1/CD133 marks stem cells and early progenitors in mouse small intestine. Gastroenterology 2009;136(7):2187–94 e1.

[29] Potten CS, Booth C, Tudor GL, Booth D, Brady G, Hurley P, Ashton G, Clarke R, Sakakibara S, Okano H. Identification of a putative intestinal stem cell and early lineage marker; musashi-1. Differentiation 2003;71(1):28–41.

[30] Saad RS, Ghorab Z, Khalifa MA, Xu M. CDX2 as a marker for intestinal differentiation: Its utility and limitations. World J Gastrointest Surg 2011;3(11):159–66.

[31] Yang Q, Bermingham NA, Finegold MJ, Zoghbi HY. Requirement of Math1 for secretory cell lineage commitment in the mouse intestine. Science 2001;294(5549):2155–8.

[32] Demitrack ES, Samuelson LC. Notch regulation of gastrointestinal stem cells. J Physiol 2016;594(17):4791–803.

[33] Grenov B, Briend A, Sangild PT, Thymann T, Rytter MH, Hother AL, Molgaard C, Michaelsen KF. Undernourished Children and Milk Lactose. Food Nutr Bull 2016;37(1):85–99.

[34] Gambarara M, Diamanti A, Ferretti F, Papadatou B, Knafelz D, Pietrobattista A, Castro M. Intractable diarrhea of infancy with congenital intestinal mucosa abnormalities: outcome of four cases. Transplant Proc 2003;35(8):3052–3.

[35] Kahvecioglu D, Yildiz D, Kilic A, Ince-Alkan B, Erdeve O, Kuloglu Z, Atasay B, Ensari A, Yilmaz R, Arsan S. A rare cause of congenital diarrhea in a Turkish newborn: tufting enteropathy. Turk J Pediatr 2014;56(4):440–3.

[36] Rehfeld JF. A centenary of gastrointestinal endocrinology. Horm Metab Res 2004;36(11-12):735–41.

[37] Latorre R, Sternini C, De Giorgio R, Greenwood-Van Meerveld B. Enteroendocrine cells: a review of their role in brain-gut communication. Neurogastroenterol Motil 2016;28(5):620–30.

[38] Wang J, Cortina G, Wu SV, Tran R, Cho JH, Tsai MJ, Bailey TJ, Jamrich M, Ament ME, Treem WR, Hill ID, Vargas JH, Gershman G, Farmer DG, Reyen L, Martin MG. Mutant neurogenin-3 in congenital malabsorptive diarrhea. N Engl J Med 2006;355(3):270–80.

[39] Lo YH, Chung E, Li Z, Wan YW, Mahe MM, Chen MS, Noah TK, Bell KN, Yalamanchili HK, Klisch TJ, Liu Z, Park JS, Shroyer NF. Transcriptional Regulation by ATOH1 and its Target SPDEF in the Intestine. Cell Mol Gastroenterol Hepatol 2017;3(1):51–71.

[40] Heijmans J, van Lidth de Jeude JF, Koo BK, Rosekrans SL, Wielenga MC, van de Wetering M, Ferrante M, Lee AS, Onderwater JJ, Paton JC, Paton AW, Mommaas AM, Kodach LL, Hardwick JC, Hommes DW, Clevers H, Muncan V, van den Brink GR. ER stress causes rapid loss of intestinal epithelial stemness through activation of the unfolded protein response. Cell Rep 2013;3(4):1128–39.

[41] Das B, Okamoto K, Rabalais J, Marchelletta RR, Barrett KE, Das S, Niwa M, Sivagnanam M. Congenital Tufting Enteropathy-Associated Mutant of Epithelial Cell Adhesion Molecule Activates the Unfolded Protein Response in a Murine Model of the Disease. Cells 2020;9(4).

[42] Huang L, Yang Y, Yang F, Liu S, Zhu Z, Lei Z, Guo J. Functions of EpCAM in physiological processes and diseases (Review). Int J Mol Med 2018;42(4):1771–85.

[43] Donowitz M, Cha B, Zachos NC, Brett CL, Sharma A, Tse CM, Li X. NHERF family and NHE3 regulation. J Physiol 2005;567(Pt 1):3–11.

[44] Dahlqvist A. Assay of intestinal disaccharidases. Anal Biochem 1968;22(1):99–107.

