## Supplemental results for "Aberrant Epithelial Differentiation Contributes to Pathogenesis in a Murine Model of Congenital Tufting Enteropathy"

Figure S1

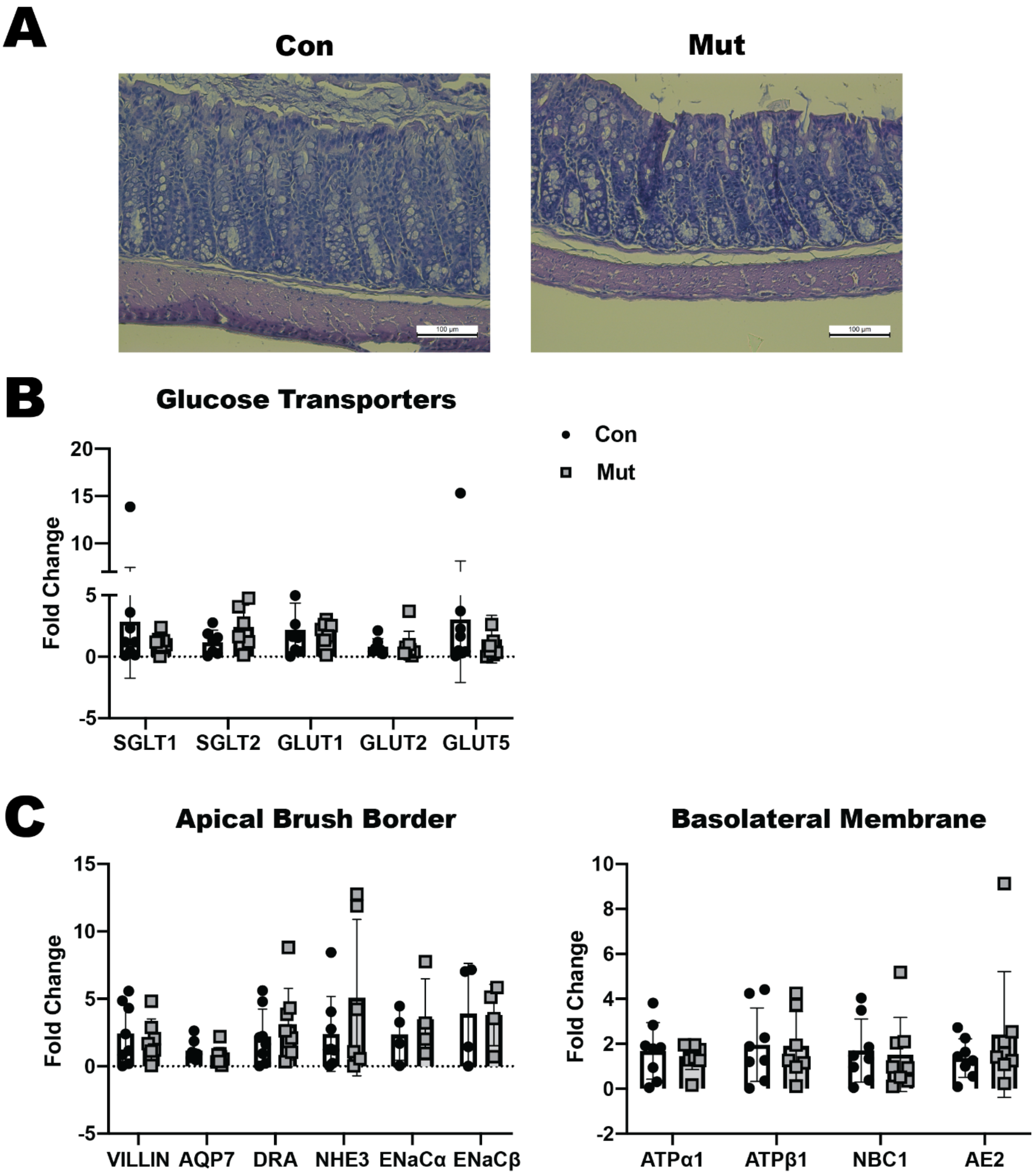


**Supplemental Figure S1: Evaluation of colonic epithelial cell markers in CTE mice (A)** Representative brightfield H&E images of colonic tissue sections from control (Con) and CTE mutant (Mut) mice (n=3). Scale bar denotes 100 $\mu$m. **(B)** mRNA expression for glucose transporters (*SGLT1, SGLT2, GLUT1, GLUT2, GLUT5*) in colonic tissue lysates from of Con and Mut mice (n=7). **(C)** mRNA expression for apical brush border markers (*VILLIN, AQP7, DRA, NHE3*) (n=7) and (*ENaC*$\alpha$ *and ENaC*$\beta$) (n=4) and basolateral membrane markers (*ATPase*$\alpha$*, ATPase*$\beta$*, NBC1, and AE2*) (n=7) in colonic tissue lysates from Con and Mut mice in colonic tissue lysates from Con and Mut mice. No statistically significant differences were found for any analyte between Con and Mut.

**Figure S2**

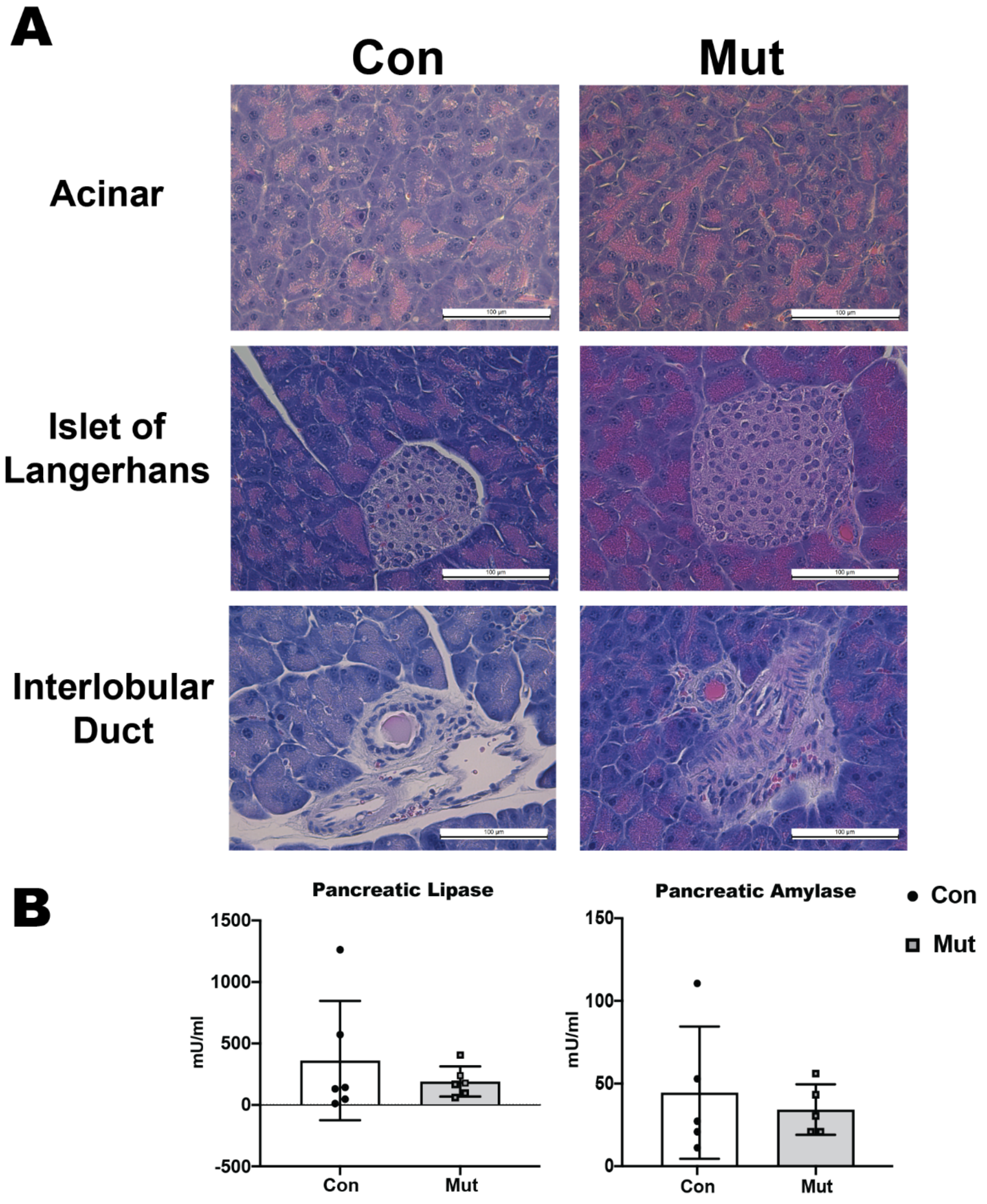


**Supplemental Figure S2 Evaluation of the pancreas in CTE mice (A)** Representative brightfield H&E images of pancreatic tissue sections showing acinar cells, islet of Langerhans, and interlobular ducts in control (Con) and CTE mutant (Mut) mice (n=3). Scale bar denotes 100 μm. **(B)** Pancreatic lipase and amylase activity (n=6) in serum from Con and Mut mice.
